# The functional and genetic associations of neuroimaging data: a toolbox

**DOI:** 10.1101/178640

**Authors:** Zhaowen Liu, Edmund T. Rolls, Jie Zhang, Ming Yang, Jingnan Du, Weikang Gong, Wei Cheng, He Wang, Kamil Ugurbil, Jianfeng Feng

## Abstract

Advances in neuroimaging and sequencing techniques provide an unprecedented opportunity to map the function of brain regions and to identify the roots of psychiatric diseases. However, the results generated by most neuroimaging studies, i.e., activated clusters/regions or functional connectivities between brain regions, frequently cannot be conveniently and systematically interpreted, rendering the biological meaning unclear. We describe a Brain Annotation Toolbox (BAT), a toolbox that helps to generate functional and genetic annotations for neuroimaging results. The toolbox can take data from brain regions identified with an atlas, or from brain regions identified as activated in tasks, or from functional connectivity links or networks of links. Then, the voxel-level functional description from the Neurosynth database and the gene expression profile from the Allen Brain Atlas are used to generate functional and genetic knowledge for such region-level data. Parametric (Fisher’s exact test) or non-parametric (permutation test) statistical tests are adopted to identify significantly related functional descriptors and genes for the neuroimaging results. The validity of the approach is demonstrated by showing that the functional and genetic annotations for specific brain regions are consistent with each other; and further the region by region functional similarity network and gene co-expression networks are highly correlated for many major brain atlases. One application of BAT is to help provide functional and genetic annotations for the newly discovered regions with unknown functions, e.g., the 97 new regions identified in the Human Connectome Project. Importantly too, this toolbox can help understand differences between patients with psychiatric disorders and controls, and this is demonstrated using data for schizophrenia and autism, for which the functional and genetic annotations for the neuroimaging data differences between patients and controls are consistent with each other and help with the interpretation of the differences.

## Introduction

Advances in non-invasive neuroimaging techniques have allowed investigation of the neural basis of human behavior^1^, ^2^ and to search for the roots of psychiatric diseases^3, 4^. Neuroimaging analysis generates results in clusters of voxels/ brain regions or in functional connectivity (FC) links between pairs of voxels or brain areas with correlated activity. The biological interpretation of these results, however, remains difficult, and we often need to look up and summarize individual studies in the literature to find biological explanations. Since each study usually has a small sample size and the results may be under powered and have a high false discovery rate^5, 6^, explanations based on these results may not be very reliable.

Recently, Neurosynth integrated results from tens of thousands of neuroimaging investigations, providing more reliable mappings between brain voxels and cognitive states than individual studies^7^. Meanwhile, the Allen Human Brain Atlas was constructed and provided a comprehensive ‘all genes-all structure’ profile of the human brain^8^. These two datasets have provided us with comprehensive knowledge for understanding the human brain at multiple scales and with multiple types or modalities of investigation. However, a huge gap still exists in using these data to interpret neuroimaging results. The mappings between voxels to function in Neurosynth, and to gene expression profiles in the Allen Brain Atlas are fine-scale (voxel-level) representations, which cannot directly provide functional or genetic meaning for brain regions consisting of clusters of voxels, or of the FCs between them. Therefore, for most neuroimaging analyses that generate results in the form of multiple brain regions or FCs, a rigorous statistical mapping from voxel-level representations (either functional or genetic) to region-level knowledge is needed.

In this research, we developed the BAT (Brain Annotation Toolbox), which, when provided with voxel-level coordinates, transfers information from Neurosynth about which functions are associated with those coordinates, and from the Allen Brain Atlas about which genes are associated with those coordinates. BAT can perform functional and genetic annotation for many neuroimaging results, either in 3D-volume space or 2D-surface space, in the form of clusters/regions or FCs. One appealing application is that BAT can provide functional and genetic descriptors for different widely used brain atlases such as Brodmann^9^, AAL2 (Automated Anatomical Labeling Atlas 2)^10^, and Craddock 200^11^. And BAT can also help identify the potential genetic and functional characteristics of newly discovered regions, such as the 97 brain regions recently identified by the Human Connectome Project (HCP), whose functional roles and genetic properties remain unclear^12, 13^. The toolbox and a user-friendly graphical user interface was developed and is publicly available at (http://www.dcs.warwick.ac.uk/∼feng/BAT).

## DATA and METHOD

### Data

#### Task activation maps

The task activation maps from the Neurosynth database (<u>http://neurosynth.org</u>) provide voxel-level functional annotation, i.e., each voxel is associated with a number of terms or tasks which help to interpret the function if that region^7^. This was obtained by integrating more than 11,000 journal articles (at the time of our analyses (May 2017) that provided the locations of task-related activations for various tasks. More than 3,000 search terms with their activation maps were obtained using text-mining techniques to analyze the abstract and automatically extract the coordinates of activations from all the articles. In our analysis, we deleted terms that were not useful in identifying tasks (e.g., ‘able’, ‘abstract’ etc.) and selected 217 terms that bear clear biological significance (details of the selection criteria are described in our previous work^14^), see Supplementary Table S1. We used forward inference maps to indicate the degree to which each voxel is consistently activated in studies that used a given term (FDR correction of P < 0.01). The activation maps were resliced to 1 × 1 × 1 mm^3^ and transformed to binary images by setting all the non-zeros entries as 1.

#### Gene expression profile

The ‘all genes-all structure’ profiles from the Allen Human Brain Atlas (AHBA) (http://human.brain-map.org/) provided the brain’s genetic expression levels for different brain regions^15^, obtained from six adult human brains from the AHBA^16^. Two of the brains were with both hemispheres and four only with the left hemisphere. The number of anatomic samples obtained from each brain varied from 363 to 946. In total, 3695 unique anatomic samples with 20,738 gene expression profiles were obtained (details of AHBA’s microarray information / data normalization: <u>http://help.brain-map.org/display/humanbrain/documentation/)</u>. To further remove individual differences and pool all the AHBA samples from different subjects together to provide voxel-level genetic knowledge, a normalization procedure was applied: for each given gene in any individual, expressions were normalized by extracting the median of the gene's expression across all samples of the individual, and were divided by the median. Then, for each AHBA tissue sample, we created a 6 mm sphere region of interest (ROI) in the MNI volume space centered on its MNI centroid coordinate. Finally, 3695 ROIs with their corresponding normalized gene expression profiles were used in our following analysis.

### Method

#### Mapping from MNI volume space to the surface space

Both of the activation maps, from Neurosynth and the gene expression in AHBA samples, were in MNI volume space (3D) and could not be directly used to interpret neuroimaging results in 2D surface space. A mapping scheme from the 3D volume space to the 2D surface space was therefore needed for both the Neurosynth and Allen Brain Atlas database. For Neurosynth, for each activation map of the 217 functional search terms, we mapped the coordinates of the activations from the MNI volume space to the Conte69 human surface-based atlas (<u>http://brainvis.wustl.edu/wiki/index.php//Caret:Atlases/Conte69_Atlas</u>) using the Human Connectome Workbench. The activation z-value of each surface vertex was transformed from the voxels in which the vertex lay. We performed this mapping for all the 217 functional terms’ activation maps in volume space, and the surface-based activation maps were obtained in the 32k Conte69 surface-based space^17, 18^.

For the Allen Brain database, we mapped the AHBA ROIs in the MNI space to the Conte69 human surface-based atlas using the same method that we used to map the activation maps. For each AHBA sample, we obtained its corresponding vertices on the surface. We manually checked the NeuroSynth activation maps and the Allen Human Brain Atlas (<u>http://atlas.brain-map.org</u>) ^15, 19^ ROIs that we mapped from their volume space to the surface space to ensure accuracy. We illustrate examples for comparison of the maps in the two spaces in Supplementary Fig 1. In the following, we use “ voxel” to denote both the 3D and 2D pixel in the brain images for convenience.

**Figure 1.**
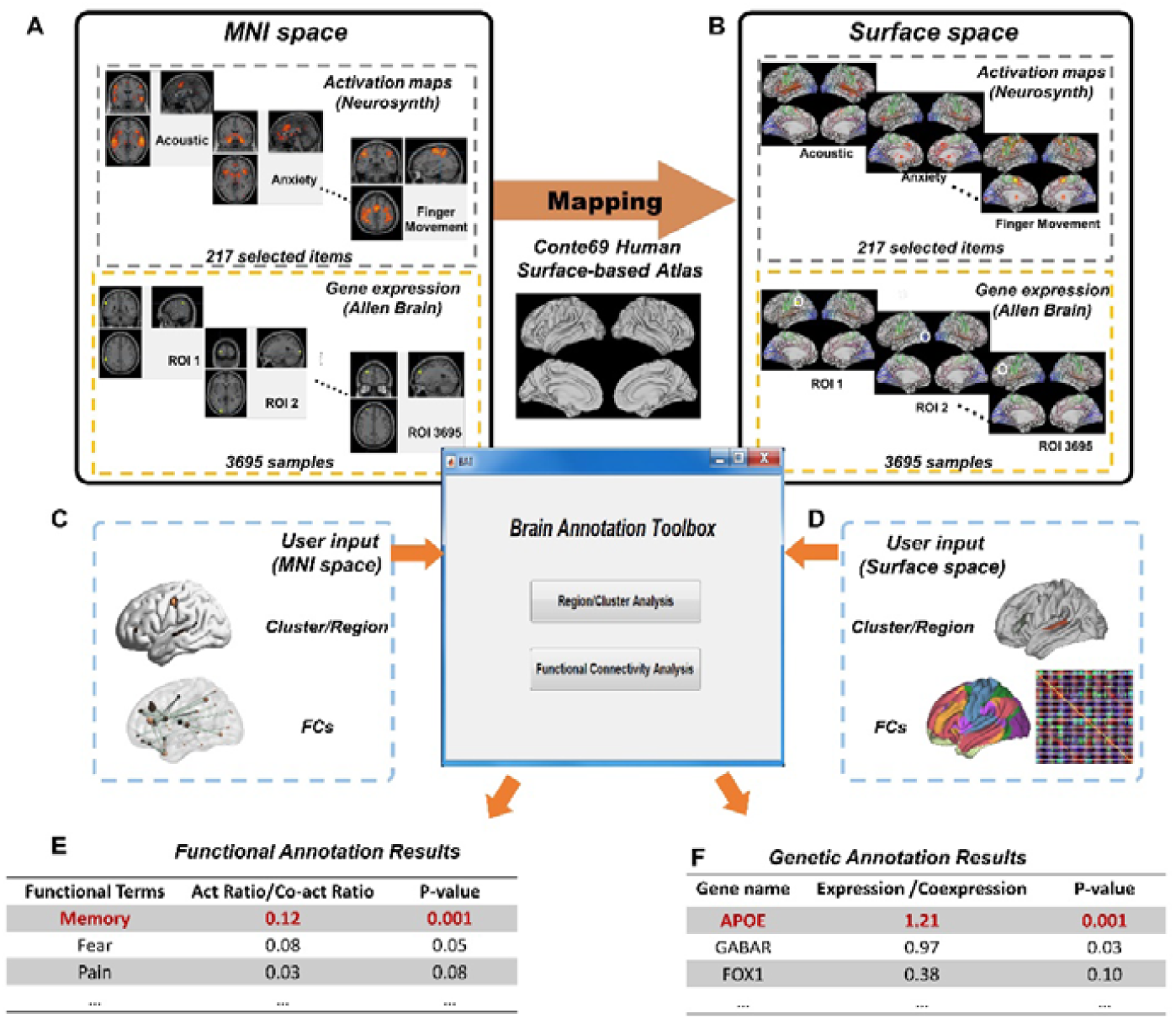
Flow chart of functional and genetic annotation analysis. (A) Upper panel: The activation maps in MNI space for the 217 functional terms from the Neurosynth database. Bottom panel: 3695 Allen Human Brain Atlas (AHBA) samples with gene expression were employed and mapped to MNI space first. (B) Upper panel: The 217 activation maps in the MNI space were then mapped to the surface-based space by registering to the Conte69 Human surface-based Atlas. Bottom panel: The 3695 ABHB samples were mapping to the Conte69 Human surface-based Atlas as well. (C, D) Two general forms of neuroimaging analysis results, i.e., clusters/regions (C) and functional connectivities (D) (either in 3D MNI space or the 2D surface space) can be analyzed by the BAT. (E) BAT can perform functional annotation analysis for user-provided neuroimaging results and provide the most-related functional terms. (F) BAT can perform genetic annotation analysis for the user-provided neuroimaging results and identify the most correlated genetic correlates.

#### Functional annotation analysis for given clusters/regions

The aim of the functional annotation analysis for clusters/regions was to provide a functional explanation or interpretation for given clusters/regions. The principle of our functional annotation analysis was the same as the widely-used gene enrichment analysis, which assumes that the co-functioning genes for the abnormal biological process underlying the study are more likely to be selected as a relevant group by high throughput screening techniques^20, 21^. Similarly, in neuroimaging research, voxels within a cluster/region have a higher probability to be co-activated by the same terms that are functionally related to the cluster/region, compared to voxels selected at random. For a given term in the Neurosynth database, the extent of activation of a given cluster/region was termed as the activation ratio (i.e., the number of activated voxels in this region divided by the total number of the voxels in the region). Further, the statistical significance of the activation was evaluated by either parametric (Fisher’s exact test) or a non-parametric approach (a permutation test performed by randomly selecting voxels within the brain background mask). In the toolbox, functional annotation analysis can be performed for a cluster/region consisting of a single component with connected voxels (e.g., a single AAL2 region), a cluster/region consisting of multiple connected components (e.g., the activated clusters obtained from a specific task), and multiple clusters/regions (e.g. multiple AAL2 regions).

For a single cluster/region, the above two kinds of statistical tests help users to infer which functional terms are significantly related to it. The parametric test is based on the Fisher’s exact test which is widely used in gene enrichment analysis^21, 22^, and the null hypothesis is that there is no relation between whether a voxel lies within a cluster/region and whether the voxel is activated for a given term. Under this null hypothesis, we can model the number of voxels in a cluster/region that are activated by a given term by the Hypergeometric distribution. Supposing there are *x* activated voxels in the cluster/region for the given term, we can get the *p*-value by simply computing the probability of observing *x* or more activated voxels in the cluster/region, see Eq.1 for details.

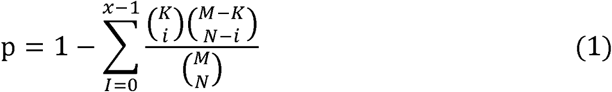

where *N* and *M* are the number of voxels in the cluster/region and the background mask, respectively; and *x* and *K* are the number of activated voxels in the cluster/region and the background.

For the statistical test based on a non-parametric permutation test, three approaches are used, differentiated by the way in which the spatial structure of the voxel in the cluster/region is considered. The first one is the most efficient and is suitable for all forms of cluster/regions. It randomly selects non-overlapping voxels within the background (with the same number as those in the given clusters/regions) and regardless of their spatial relationship. The second is suitable for the clusters/regions consisting of a single spatially connected component. For example, to annotate a region in the AAL template, we select the same number of voxels as that in the given region and these voxels are also spatially adjacent in the background. The third is for the clusters/regions consisting of multiple spatially connected components. In this case, we randomly select non-overlapping connected components (with the same number as that in the given cluster/region), each consisting of spatially adjacent voxels (and with the same number as those in the components in the given cluster/region) from the background. After determining the voxel/vortex selection approach, BAT runs the permutation *N* times (*N* is the number of permutations defined by the user), to get a null distribution of the activation ratio for each term. The observed activation ratio is then compared with the null distribution to get the corresponding p-value.

#### Genetic analysis for the clusters/regions

Based on the gene expression data from AHBA, the BAT’s genetic analysis for the clusters/regions can provide the whole genomic gene expression profiles for the cluster/regions of interest, and help to identify the differentially expressed genes. The details for our genetic annotation analysis for clusters/regions are as follows.

First, with a given background mask, we retain AHBA samples with more than 50% of voxels that are also present in the background mask to perform further analysis (we term these samples as the background AHBA samples). Then, for each background AHBA sample, we map it to one of the given clusters/regions, that which has the largest number of overlapping voxels with this AHBA sample. The gene expression profile of each region/cluster is defined as the average gene expression of all the samples mapped to the cluster/region. We then adopt permutation analysis to identify the differentially expressed genes in the given clusters/regions (compared with all samples in the background). Two methods are used for sample selection in the background: 1. randomly selected AHBA samples from the background without repetition, and 2. randomly selected AHBA samples in the background samples but not the ones that were already mapped to the region/ROI. Then for each cluster/region in each permutation run, we randomly select the same number of AHBA samples as those that are mapped to the cluster and calculate the average gene expression profiles across all selected samples. A null distribution for each gene was thereby obtained, allowing us to rank each gene in its null distribution and got its corresponding p-value for over-expression or down-expression.

#### Functional annotation analysis for functional connectivity (FC)

The BAT can also perform functional enrichment analysis for a FC or set of FCs constituting a network. A difference from previous analyses described for the BAT is that now the input data consist of a set of significant functional connectivity (FC) links. For example, we can determine the functions associated with the underlying FCs/networks identified by either a ROI-based approach or a brain-wide association study (BWAS). This is especially useful for the altered FCs identified in case-control studies. At the outset, we make it clear that functional connectivities measured as correlations between brain areas are not being computed in the BAT. Instead, we make use of the evidence that the FC between two nodes even with resting state fMRI is typically correlated with the activation in a particular task of these two nodes ^23^-^27^, and it is the latter that the ‘functional connectivity’ analyses in the BAT reflect, as follows.

An image map for the regions that were connected by the FCs and a list of all the FCs of interest are required to perform the analysis. First, to measure to what degree two regions connected by a FC are co-activated in a certain term, or task, we defined the “ co-activation ratio” as the average proportion of activated voxels of these two regions for a FC. If one of the two regions had no voxel activated in the activation map for the function term, we set the co-activation ratio of the FC as 0. For a functional network consisting of multiple FCs, its extent of activation is defined as the mean co-activation ratio, i.e., the average of the co-activation ratio of all the FCs in the network. In calculating what is described in this paper as ‘functional connectivity’, the activity of a node (i.e. a region of interest such as an AAL2 area) for a particular search term was calculated by the mean activation of the voxels in that node in that task. If in an analysis involving multiple FC links some nodes appear *n* times, then the activity of that node is weighted by the number *n* of such links so that its annotations contribute in this proportion to the annotations for this set of ‘functional connectivity links’.

Further, the significance of the network’s mean co-activation ratio is assessed using non-parametric permutation tests. Two methods for randomly selecting the regions connected by the FCs are used. The first is suitable for a brain network consisting of a moderate number of FCs (e.g., less than 20) and in which the brain regions connected by the FCs only occupy a small fraction of the brain (so that we can randomly select the same number of non-overlapping regions from the background). Using this method, in each permutation run, BAT randomly selects the same number of non-overlapping regions consisting of the same number of adjacent voxels as those in the resulting list from the background. The second method is suitable for FCs that connect regions from whole brain atlases, e.g. the FCs obtained from regional-level brain-wide association analysis which produce a network with a large number of FCs that cover much of the brain. In such a situation, it is not feasible to randomly select the same number of non-overlapping regions from the background. We then randomly select the same number of regions as those in the FC list from the whole brain atlas being used. Given the permutation method, the mean co-activation ratio of the FCs for each of the functional terms can be calculated based on the randomly selected regions. The null distribution of the mean co-activation ratio of the FCs for each of the functional terms are constructed after running the permutation multiple times. Based on the null distribution of a functional term, we can obtain a p-value for our observed mean co-activation ratio as the proportion of permutations in which with the randomly produced mean co-activation ratio is larger than the observed mean co-activation ratio.

#### Genetic analysis for the FCs

BAT can also identify genetic correlates for the given FCs, e.g., finding genes that might regulate the functional co-activation between two brain regions. First, the gene expression profile for each region involved in the given FCs is obtained (the same as for the ‘Genetic analysis for the clusters/regions’). For each FC, the co-expression value of a gene is defined as the outer product of its expression in these two regions^28^. Then, for each gene, we can obtain an average co-expression value, i.e., the mean of the gene’s co-expression for all FCs. Permutation analysis was applied to estimate the significance of the average co-expression value for each gene: first, in each permutation run, for each region in the FC list, we randomly select the same number of AHBA samples from the background as those mapped to the regions of the given FCs without repetition, and calculate a new gene expression profile for the region, based on which we can obtain the average co-expression values for each gene. A p-value for the real average co-expression value was obtained for each gene.

## RESULTS

### The implementation of BAT

BAT is implemented as a free and open-source Matlab toolbox. The toolbox provides simple commands for users to perform genetic and functional annotation analysis on clusters/regions and FC results. A graphical user interface (GUI) is provided for users to perform the annotation analysis. A visual interface is also implemented to provide 3-D interactive visualization for the annotation results.

BAT provides a flexible setting so that users can choose to meet their requirements. BAT comes with a User Manual to describe its use. Before analysis, a background mask needs to be specified, which is a binary image describing the areas in which the user wishes to perform their analysis for permutation, e.g. the whole brain, cerebral cortex, subcortical areas, or a specific region. The user can choose whether or not to perform permutation (and to specify the permutation method and number of permutations to use). The default settings of the BAT are given in Supplementary Table S2.

### Functional and genetic annotation for well-known brain atlases

Using BAT, we performed functional and genetic annotation analysis for several widely-known brain atlases, including the Brodmann^9^, AAL2 (Automated Anatomical Labeling Atlas 2)^10^, the new Human Connectome Project (HCP) atlas, and Craddock 200^11^, as detailed in Supplementary Table S3.

In particular, we highlight here the annotation results for Brodmann areas. We manually compared the functional annotation for 32 Brodmann areas (with significant annotation results, i.e., the region had at least one significant functional annotation by permutation test, p< 0.05) with those summarized in Wikipedia (wiki) (*https://en.wikipedia.org/*), to validate our approach. The annotations for all 32 regions provided by BAT were in agreement with those in Wikipedia, i.e. there was a large extent of overlap between the functions we identified in these regions and those described in Wikipedia, see Supplementary Table S4. The annotation results for other atlases can be found at our website (http://www.dcs.warwick.ac.uk/∼feng/BAT). The functional and genetic annotations provided by BAT provide a valuable complement to these widely-used atlases.

### Functional and genetic annotation for the new brain atlas from HCP

In addition to traditional brain atlas, we also applied BAT to the recent HCP (The Human Connectome Project) Brain Atlas^12^. Using multi-modal data from the HCP, each hemisphere of the human cerebral cortex was parcellated into 180 different cortical areas. Among the 180 areas, 83 are consistent with previous reports, and 97 were newly identified in the HCP. This was an important advance, but did not address the genetic features underlying the 180 cortical areas, nor in detail the functions of each of the cortical areas^13^.

To illustrate the information that BAT makes available for the 180 cortical areas in the HCP Brain Atlas, we describe the results for two selected areas: one is the hippocampus, and the other is a cortical area newly identified with the HCP Brain Atlas, the ‘Middle Insular Area’ (MI). As the functional and genetic annotations for the two regions are all available for the left hemisphere, here we focus on the left Hippocampus and MI, with details in Figure 2.

**Figure 2.**
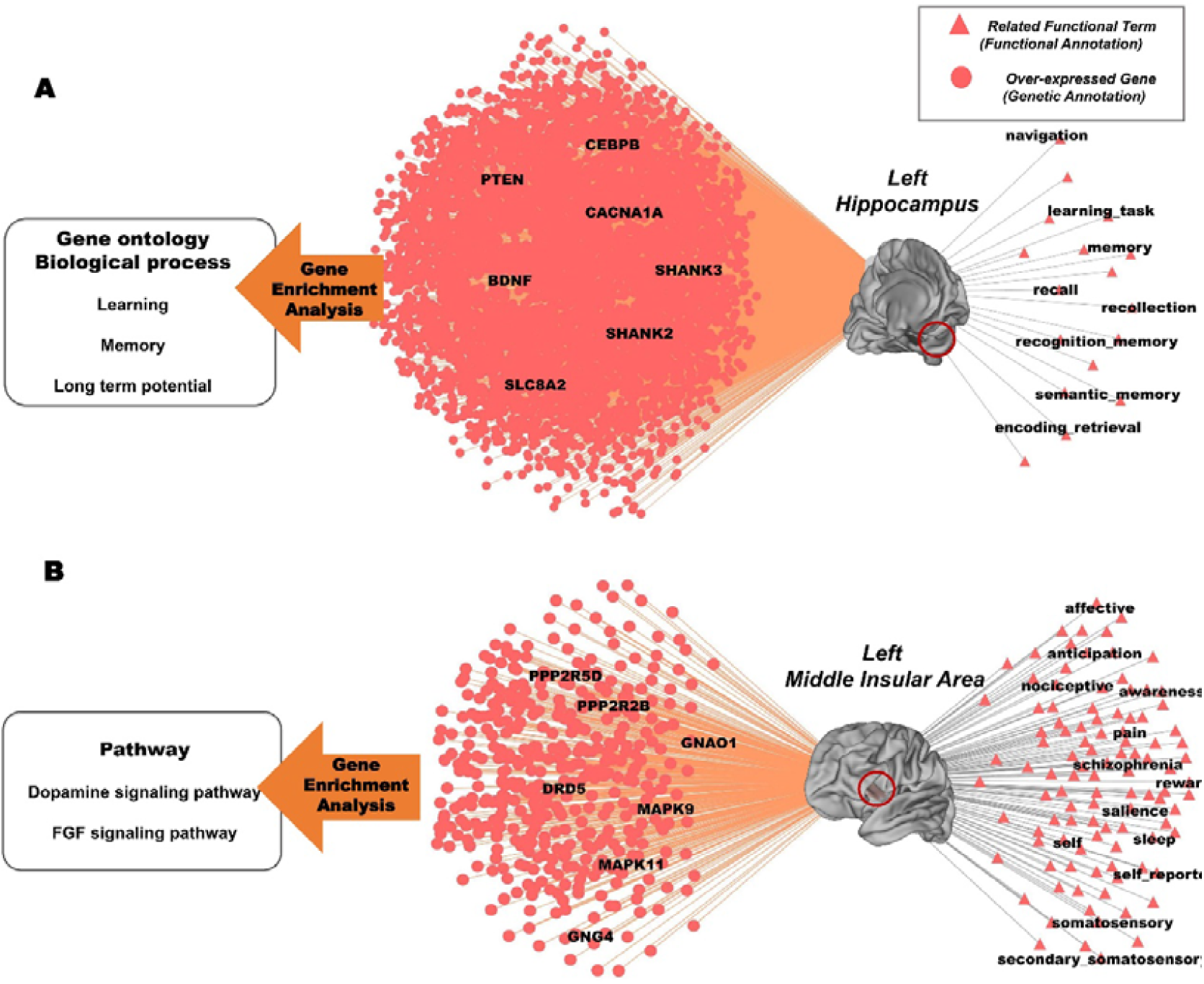
Illustration of the functional and genetic annotations of two cortical areas in the Human Connectome Project (HCP) Brain Atlas. (A) Left Hippocampus: seventeen functional terms, including memory-related ones such as ‘memory’, ‘recognition memory’,’ Semantic memory’, were found to be significantly associated with the left hippocampus (p< 0.05). For genes, 4839 genes were found to be overexpressed including the BDNF. Gene enrichment analysis shows that these genes are enriched in memory and learning related Gene Ontology (GO) biological processes such as ‘Learning’, ‘Memory’ and ‘Long term potentiation’. (B) Left Middle Insular (MI) Area: 105 functional terms were found to be significantly related to the MI area (p< 0.05), ‘affective’, ‘awareness’ ‘reward’, ‘self’, ‘salience’, ‘pain’ ‘schizophrenia’, ‘somatosensory’ are among the 12 that can survive the Bonferroni correction. 415 genes were over-expressed in the MI area and enriched in the Dopamine signaling pathway and FGF signaling pathway.

For the Hippocampus, 17 out of 217 functional terms, including ‘memory’, ‘episodic memory’, ‘navigation’, ‘recall’, ‘learning task’ etc, were found to be significantly associated with the hippocampus (p< 0.05, permutation test) (Fig. 2 and Supplementary Table S5). For genes, 4839 genes were found to be significantly overexpressed (i.e. genes expressed in this brain region or cluster or clusters more than in the rest of the brain) (p< 0.05, Bonferroni corrected). Gene enrichment analysis of these genes (using the software Toppgene^29^) revealed that processes such as ‘learning or memory’ (p=2.77e-7), ‘learning’ (p=2.16e-5) and ‘memory’ (p=3.68e-5) are significantly associated genes. The biological gene pathway “ long-term potentiation” underlying learning and memory was also found to be significantly enriched. These genes are also related to abnormal mouse phenotypes, such as ‘abnormal synaptic transmission’, ‘abnormal long term potentiation’ and ‘abnormal synaptic plasticity’.

Next, we summarize the results for a newly discovered cortical area, the MI, which is part of the insular cortex. BAT identified 105 out of 217 functional terms that were significantly related to activations produced in the MI area (p< 0.05, permutation test). Among the 105 functional terms, 12 could survive Bonferroni correction, including ‘affective’, ‘awareness’, ‘reward’, ‘self’, ‘salience’, ‘pain’, ‘schizophrenia’, ‘somatosensory’ and so on. For genes, we found that 415 genes were significantly over-expressed in the MI area (p< 0.05, Bonferroni corrected), significantly enriched in pathways that included the ‘dopamine signaling pathway’ (p=5.98e-6) and ‘FGF signaling pathway’ (p=2.131e-5). Interestingly, almost all the functional terms identified above were related to the dopamine pathway, the same as in the genetic annotation, suggesting consistency between the functional and genetic annotation, and thus verifying the usefulness of our approach. Detailed results for these two regions are provided in Supplementary Table S5.

### Functional and genetic annotations for abnormal clusters identified in Autism

To illustrate how BAT can help to gain insight into the biological meaning of neuroimaging results, we performed a functional and genetic annotation analysis for the clusters obtained in a brain-wide association analysis (BWAS) of functional connectivity for autism^30^, in which a statistical map is obtained by meta-analysis (with the Liptak-Stouffer Z-score approach) that integrates BWAS results from 16 imaging sites (418 patients and 509 controls). Then, Gaussian random field correction (cluster defining threshold: absolute Z=5.5, cluster size p< 0.05) was performed and 23 clusters consisting of voxels that had significant functional connectivity changes were obtained.

We then fed these clusters to BAT, and found they are functionally enriched in ‘autism’ and autism-related functional terms including ‘communication’, ‘self’, ‘social’, ‘theory of mind’ etc. For genetic analysis, 1117 genes were found to be significantly over-expressed in the above clusters (p< 0.05, Bonferroni corrected), which were also significantly enriched in ‘autism’ (q=1.17e-07, Bonferroni corrected) and biological processes closely related to autism, such as ‘synaptic signaling’^31^, ‘neurogenesis’^32^ etc. Interestingly, these clusters were functionally and genetically enriched in several other psychiatric diseases such as schizophrenia and depression, indicating common genetic factors underlying these mental disorders^33^, detailed in Supplementary Table S7. All the above functional and genetic annotation results are summarized in Figure 3.

**Figure 3.**
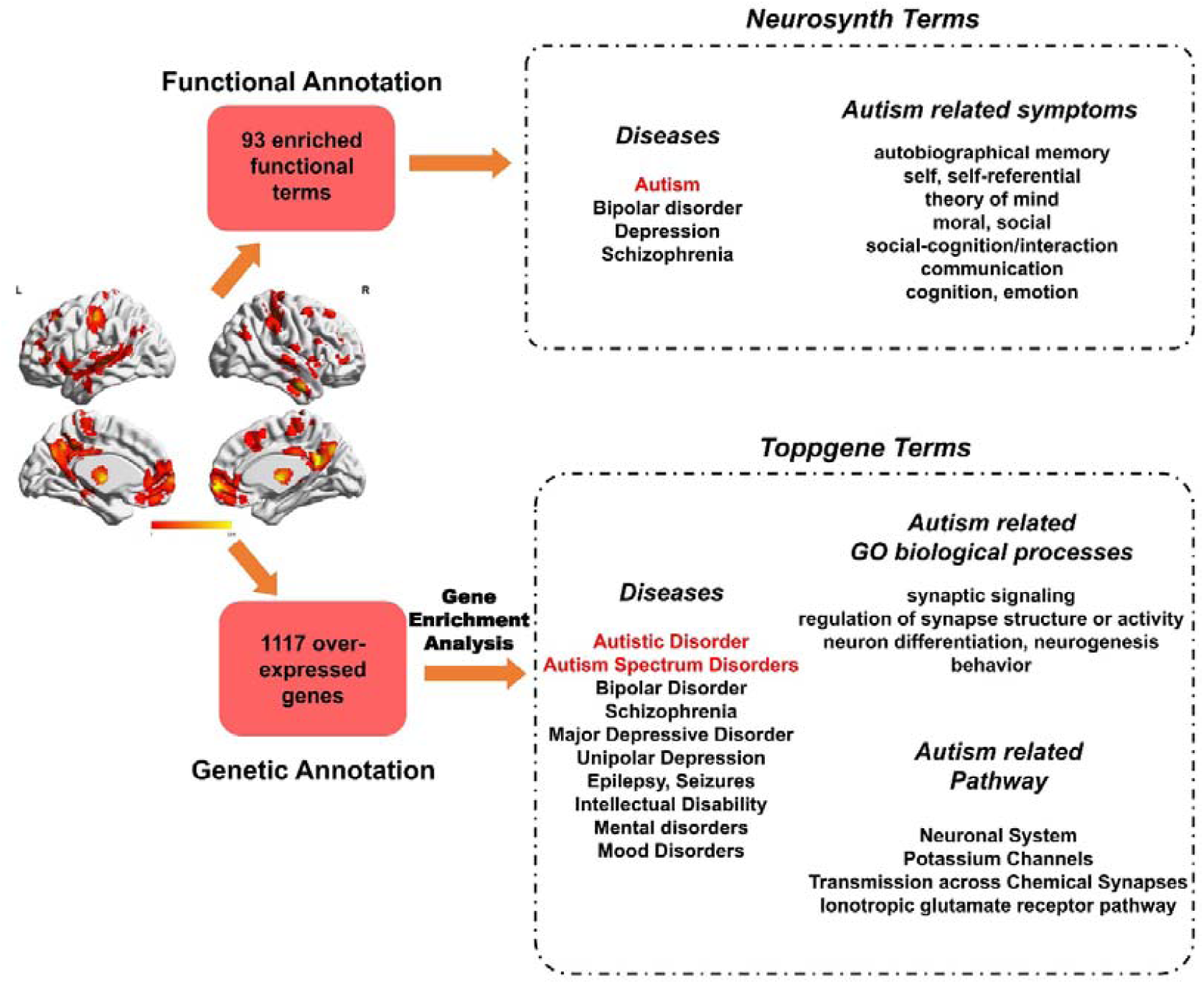
The functional and genetic annotation for clusters obtained from the Autism BWAS results. 83 functional terms were found to be significantly related to the clusters, including ‘Autism’ and several Autism-related symptoms such as ‘autobiographical memory’, ‘communication’, ‘self-referential’, ‘theory of mind’ and so on. Several Neurosynth terms for mental diseases, e.g. ‘Bipolar disorder’,’ Schizophrenia’ and ‘Depression’ were also found to be significant. For genetic analysis, 1117 genes were identified to be over-expressed, which are also functionally enriched in the disease terms ‘Autistic Disorder’ and ‘Autism Spectrum Disorders’, and several autism-related GO biological processes and pathways. The gene enrichment analysis was performed using the Toppgene software.

### Functional and genetic annotations for altered functional connectivities and networks in schizophrenia

To illustrate BAT’s capability in helping to analyze neuroimaging results in the form of functional connectivity (or a brain network defined by a set of FCs), we further used BAT to perform functional and genetic analysis on the significantly different functional connectivity links identified in chronic schizophrenia patients^34^. A resting-state brain-wide functional connectivity analysis was performed on multiple sites (with a total of 789 participants including 360 patients)^34^, and the results were integrated by meta-analysis. We performed BAT on the 89 FCs that were significantly increased in chronic schizophrenia compared to controls.

We found that this dysregulated network of 89 FCs is significantly enriched in 43 functional terms (permutation test, p< 0.05), including ‘schizophrenia’ (p=0.0349) and ‘hallucination’ (p=0.0081). Interestingly, these significantly increased FCs were also found to be significantly correlated with hallucination^34^, which is an item in the Positive subscale of the PANSS score. In addition, several other terms related to cognitive processes were also found to be significantly enriched, including “ attention” and “ memory”, detailed in Supplementary Table S8. These cognitive functions are known to be impaired in patients with schizophrenia^35, 36^. Finally, of all the identified functional terms, “ sleep” was the most significant (p< 1e-4). Disturbed sleep is frequently encountered in patients with schizophrenia and is an important part of its pathophysiology^37^.

For the genetic analysis, we selected those FCs whose associated brain regions had more than 5 AHBA samples, and this left 47 of the 89 FCs for genetic analysis. In total, 1523 genes were identified to be significantly co-expressed (p< 0.05, Bonferroni corrected) in the regions connected by these 47 FCs. These genes were significantly enriched in biological terms such as “ brain development” (p=2.987e-9), and “ neurogenesis” (p=2.258e-8, which are known to underlie the pathology of schizophrenia. Importantly, these genes were significantly enriched in the disease term “ schizophrenia” (Bonferroni correction, p=8.892e-7), and were enriched in the mouse phenotypes involving ‘abnormal sleep behavior’ (p=3.352e-3), ‘sleep disorders’ (p=5.635e-4) and ‘sleep disturbances’ (p=1.973e-3), see Figure 4.

**Figure 4.**
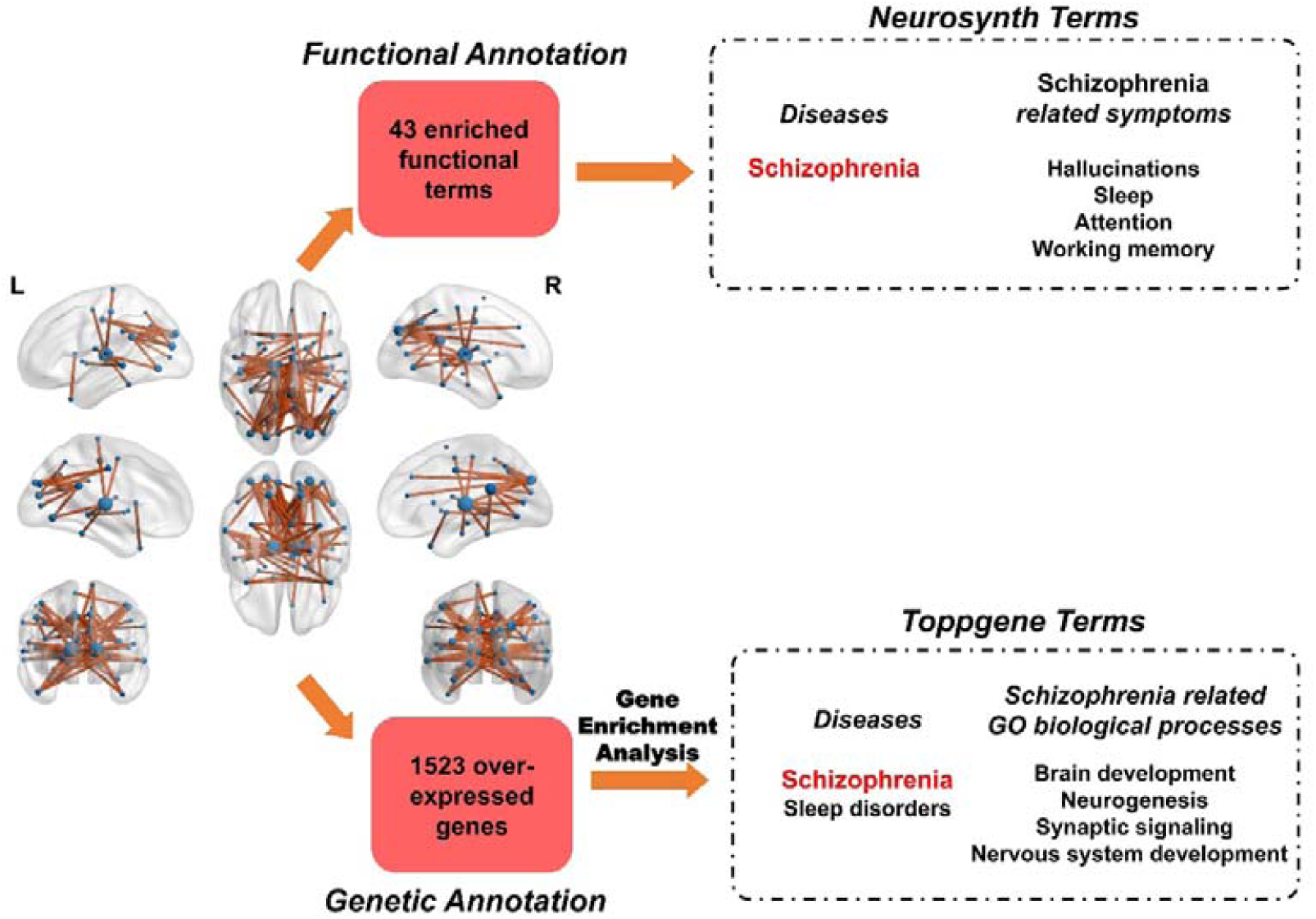
Functional and genetic annotation results for the significantly increased functional connectivity identified from chronic schizophrenia. The 89 increased FCs are significantly enriched in 43 functional terms including ‘schizophrenia’ and ‘hallucination’, “ attention” and “ memory”. 1523 genes were identified to be significantly co-expressed in the regions connected by these FCs. These genes were significantly enriched in biological terms such as “ brain development” and “ neurogenesis”.

In summary, the functional and genetic terms identified from the dysregulated network were both cross-validated, and highly consistent with the current understanding of schizophrenia, providing further evidence for the validity of the approach described here.

### Discussion

Advanced neuroimaging techniques such as fMRI have generated gigantic neuroimaging data crucial for understanding the neural basis of behavior and for exploring the pathology of psychiatric disease. However, the results obtained in neuroimaging analysis, usually in the forms of clusters of voxels/ brain regions or functional connectivities / networks, often remain hard to explain. In this research, we presented a toolbox that can provide functional and genetic annotations for brain atlas or neuroimaging results in the form of activation maps or functional connectivity, which is expected to shed insights into the biological meaning underlying these results.

In the field of bioinformatics, such an annotation analysis, gene functional enrichment analysis has already been employed to systematically dissect large ‘interesting’ gene lists from the high-throughput studies, and furthermore identify the most relevant biological processes^21^, based on the large amount of biological knowledge accumulated in public databases, i.e. Gene Ontology. During the past decades, hundreds of gene functional enrichment analysis tools have been developed and employed by tens of thousands of high-throughput studies, providing valuable insights into the underlying biological meaning of the gene analysis results.

In sharp contrast, in the neuroimaging field, large databases such as Neurosynth^7^ and AHBA^16^, have only recently been developed to provide functional / genetic knowledge for the human brain at the voxel level. However, tools for “ enrichment analysis” of neuroimaging results are still lacking. Inspired by gene enrichment analysis, we developed the BAT toolbox, which employs brain voxel-level functional and genetic knowledge to help systemically explore the region-level neuroimaging results (i.e. clusters / regions, or FCs).

BAT provides a novel method to harness the data from the Neurosynth and AHBA to perform functional and genetic annotation analysis for clusters/regions and FCs results, respectively. A user-friendly Matlab GUI and 3-D visual interface are also provided for users’ convenience. We present four examples (for clusters/regions and FCs) in the Results to illustrate the reliability of our annotation approach and to illustrate how to use BAT to search for the underlying biological meaning of the real neuroimaging results. It is noted that “ Neurosynth” also employed AHBA to identify the molecules that may participate in specific psychological or cognitive processes (“ Neurosynth-Gene”: http://neurosynth.org/genes/)^38^. However, it differs significantly from our approach in the following aspects: 1. The goal of “ Neurosynth-Gene” is to map individual cognitive phenomena to molecular processes, while the goal of BAT is to provide functional and genetic annotations for extensive neuroimaging results not necessarily confined to cognitive processes, e.g., from case-control studies. 2. BAT can provide functional and genetic annotations and corresponding p values for neuroimaging results in the form of functional connectivity or networks generated by whole-brain network analysis, which is widely used in the neuroimaging communities. This is not provided by “ Neurosynth-Gene”.

One attractive function of BAT is to help explore the newly discovered regions identified by neuroimaging technology, with unknown functions and genetic basis. We use the new parcellation of the human cortex provided by HCP as an example^12^. The 180 cortical areas in the parcellation are distinguished by multi-modal data including anatomical measurements, task-related functional magnetic resonance imaging (fMRI) of 7 tasks, and resting-state functional connectivity in a subject cohort of 210 healthy young adults. This parcellation for the human cortex is at the highest resolution to date, but neither the function nor the genetic characterization of the 180 regions, especially for the 97 newly discovery regions, are clearly known. BAT can partly solve the problem: it can provide a complementary functional and genetic interpretation for the parcellation, and researchers using the new brain parcellation in their studies can use BAT to help explore the biological meaning of their results.

We now explain why functional and genetic annotations contain similar items for a number of brain regions. Previous investigations have identified the similarity between the gene co-expression network and resting-state functional network across regions, suggesting that the functional brain network is underpinned by the gene co-expression network^39, 40^. To further validate our functional and genetic annotation we used regions selected from the Brodmann, HCP, AAL2 and Cradock atlases and computed similarity matrices between all pairs of regions for the genetic and for the functional annotations. We found that these two similarity matrices corresponded significantly, as described next. We compared the following two networks: region by region coactivation networks, and region by region gene co-expression networks, for a given brain atlas. The former was constructed by calculating the Pearson correlation coefficient between the activation ratios (of all 217 search terms or tasks) for each pair of brain regions; and the latter was obtained by calculating the Pearson correlation between the gene expression profile for each pair of brain regions. We found that the functional and genetic similarity matrices were significantly correlated, and this was found for all the brain atlases (see Figure 5; AAL2: r=0.310, p=2.9947e-78; BA r=0.4229, p=2.87e-30; CRAD r=0.2715, p=7.90e-121; HCP r=0.2635, p= 7.44e-78) adopted in this work, indicating that two brain regions with similar genetic expression profiles are more likely to have similar activation patterns. BAT has a few limitations. Currently, the functional annotation analysis of BAT is based on the 217 selected functional terms for Neurosynth, which cannot capture all the functional terms associated with all brain areas. For the genetic annotation analysis, the samples from the AHBA do not cover the whole brain. Therefore, for regions/clusters or FCs that do not have enough AHBA samples (e.g. less than 5 samples) mapped to them, genetic analysis is not possible. Further effort should involve integrating activation maps from all available meta-analysis databases (such as Brainmap^41^) and gene expression profiles (such as that from Gene Expression Omnibus^42^), to provide a more comprehensive and reliable functional and genetic annotation for neuroimaging analysis. An advantage of BAT is that the Matlab source code is provided with the toolbox, allowing users to understand what is being computed, and to enable users to develop further enhancements.

**Figure 5.**
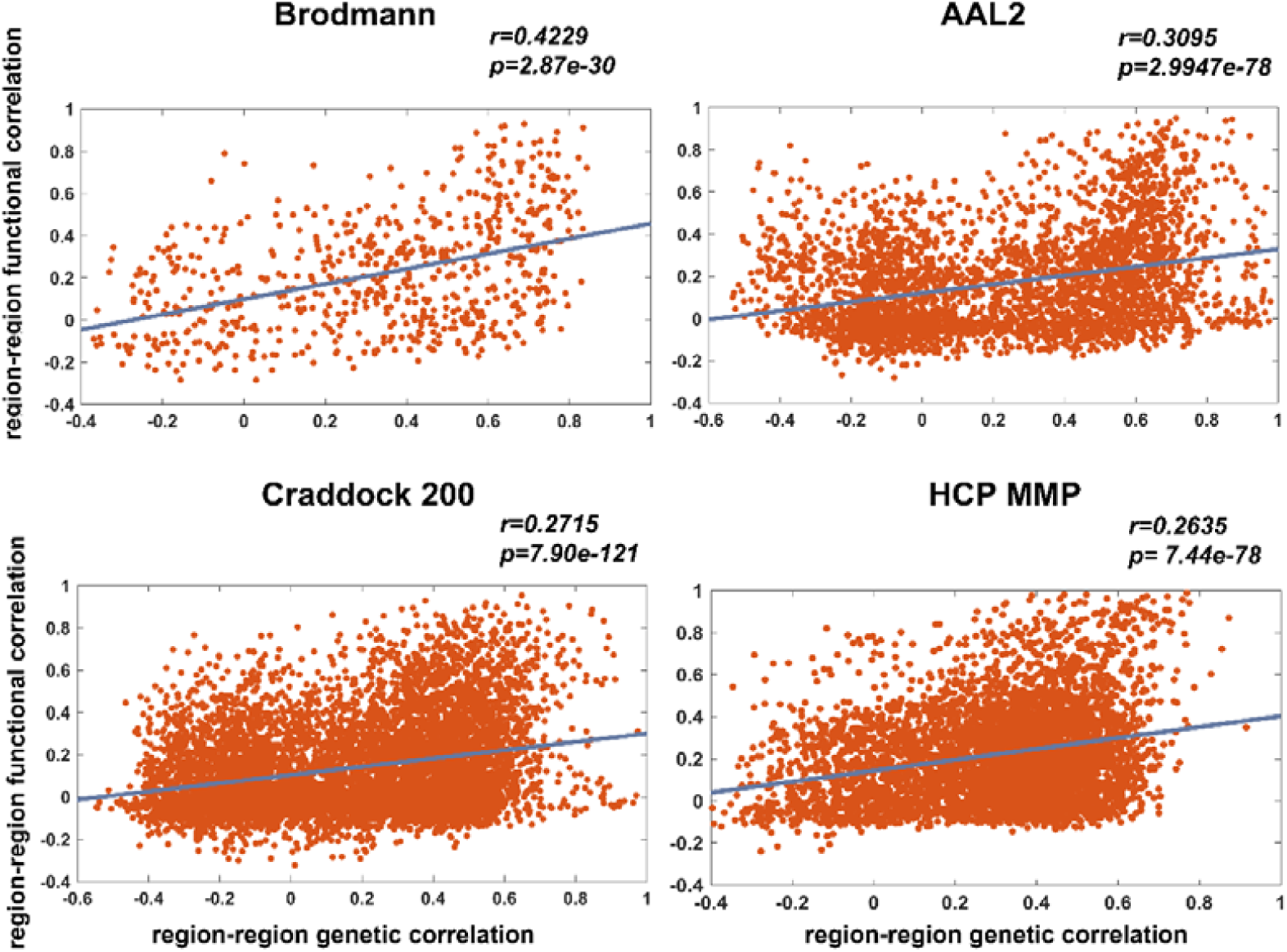
A high correlation was found between the region by region co-activation network, and the region by region gene co-expression network for A. the Brodmann atlas, B. the AAL2 atlas, C. the Craddock atlas, D. the HCP atlas. Each dot in the figure represents an edge in the region by region network. The coactivation network was obtained by calculating the correlation coefficient between the activation ratios (of all 217 terms or tasks) for each pair of brain regions in a given atlas, and the gene co-expression was obtained by calculating the correlation between the gene expression profile for each pair of brain regions in the same atlas.

